# Proximal dendritic localization of NALCN channels underlies tonic and burst firing in nigral dopaminergic neurons

**DOI:** 10.1101/2022.07.17.500361

**Authors:** Suyun Hahn, Ki Bum Um, Hyun Jin Kim, Myoung Kyu Park

**Affiliations:** Department of Physiology, Sungkyunkwan University School of Medicine, 2066, Seoburo, Jangan-gu, Suwon, 16419, Korea; Samsung Biomedical Research Institute, Samsung Medical Center, 81 Irwon-Ro Gangnam-gu, Seoul, 06351, Korea

**Keywords:** Dopaminergic neurons, Substantia nigra, Pacemaker channels, Pacemaking, TRPC3, NALCN, Ion channels, Burst firing, Tonic firing

## Abstract

In multipolar nigral dopamine (DA) neurons, the highly excitable proximal dendritic compartments (PDCs) and two Na^+^-permeable leak channels, TRPC3 and NALCN, play a key role in pacemaking. However, the causal link between them is unknown. Here we report that the proximal dendritic localization of NALCN underlies pacemaking and burst firing in DA neurons.

Our morphological analysis of nigral DA neurons reveals that TRPC3 is ubiquitously expressed in the whole somatodendritic compartment, but NALCN is localized within the PDCs. Blocking either TRPC3 or NALCN channels abolished pacemaking. However, only blocking NALCN, not TRPC3, degraded burst discharges. Furthermore, local glutamate uncaging readily induced burst discharges within the PDCs, compared with other parts of the neuron, and NALCN channel inhibition dissipated burst generation, indicating the importance of NALCN to the high excitability of PDCs. Therefore, we conclude that PDCs serve as a common base for tonic and burst firing in nigral DA neurons.

## Introduction

Midbrain dopamine (DA) neurons are slow pacemakers that basically generate regular tonic firing, but strong afferent synaptic inputs can evoke high-frequency phasic or burst discharges (Gonon, 1988; Grace et al., 2007, Blythe et al., 2009). While tonic firing of midbrain DA neurons maintains ambient DA levels in the target areas, burst firing triggers DA surges (Morikawa and Paladini, 2011). DA is a key neurotransmitter that is critically important for incentive motivation and reinforcement learning (Schultz, 2007; Wise, 2004). In addition, aberrant excitabilities in DA neurons underlie many kinds of neuropsychiatric diseases, including Parkinson’s disease (Surmeier et al., 2012). Therefore, ion channel mechanisms for pacemaking and burst firing have been a longstanding research topic in DA neurons. Ion channels and morphological features are the two major factors that determine the rate and pattern of neuronal firing.

Midbrain DA neurons extend 3–6 primary dendrites, and interestingly, their axons most often originate in one of several proximal dendritic compartments (PDCs) (Tepper et al., 1987; Häusser et al., 1995; Gentet and Williams, 2007). An axon initial segment (AIS) is a site at which action potentials (APs) first arise (Gentet and Williams, 2007; Kole et al., 2008), but both tonic and burst firing can be generated in the absence of an axon without significant alterations in spontaneous firing rates (Kim et al., 2004; Kim et al., 2007; Jang et al., 2014; Kimm et al., 2015). Therefore, it could be the soma and dendrites that formulate the firing rate and patterns in DA neurons, even though the AIS might affect the AP threshold and shape the final outputs of firing (Meza et al., 2018). In addition, multipolar DA neurons produce synchronized spontaneous APs throughout the entire somatodendritic compartment according to the pacemaking cycle (Wilson & Callaway, 2000; Guzman et al. 2009). Therefore, the tight electrical coupling between the soma and excitable dendritic compartments underlies the pacemaking mechanism of midbrain DA neurons (Wilson & Callaway, 2000; Guzman et al. 2009). Assuming that intrinsic excitability is similar across the entire somatodendritic membrane, dendritic compartments with a high surface area/volume ratio are faster oscillators than the soma, so they play an accelerating role in pacemaking (Wilson & Callaway, 2000; Jang et al., 2014). We previously reported that PDCs, not the whole dendritic compartment, mainly participate in the pacemaking of nigral DA neurons (Jang et al., 2014). Furthermore, we showed that PDCs are more excitable than the soma and distal dendritic compartments, suggesting that PDCs dominantly drive pacemaking in nigral DA neurons (Jang et al., 2014).

Very recently, we reported that two nonselective cation channels, the sodium leak channel (NALCN) and transient receptor potential-canonical 3 (TRPC3) channel, produce a sustained leak-like inward current that is responsible for the slow and regular pacemaking of nigral DA neurons (Um et al., 2021). NALCN is a Na^+^-permeable leak channel that produces background Na^+^-leak conductance and determines the resting membrane potential and excitability of many central neurons (Lu et al., 2007; Lutas et al., 2016; Shi et al., 2016; Hahn et al., 2020). TRPC3 is another leak channel that opens constitutively, leading to slow depolarization in many neurons (Dietrich et al., 2003; Zhou et al., 2008; Um et al., 2021). Although NALCN and TRPC3, as subthreshold leak channels, underlie the slow depolarization responsible for pacemaker activity in nigral DA neurons (Um et al., 2021), little is known about the relationships between these two pacemaker ion channels and the highly excitable PDCs (Jang et al., 2014). Furthermore, pacemaker DA neurons can generate high-frequency burst discharges in response to primary rewards or cues that predict rewards via the glutamatergic synapses (Schultz et al., 1998; Morikawa and Paladini, 2011). Although burst generation in midbrain DA neurons is known to require N-methyl-D-aspartate receptors (NMDARs) in dendritic compartments (Deister et al., 2009; Overton and Clark, 1992; Seutin et al., 1993), strong glutamatergic stimuli could evoke burst firing by means of the intrinsic excitability of dendritic compartments (Deister et al., 2009). Therefore, the NALCN and TRPC3 channels could contribute to the intrinsic excitability of dendrites and participate in the burst firing of DA neurons. In this work, we report that functional TRPC3 channels exist in the whole somatodendritic compartment, but NALCN channels are highly localized in PDCs in nigral DA neurons. The proximal dendritic localization of NALCN channels imparts the high excitability associated with PDCs. Therefore, multiple highly excitable PDCs appear to serve as a common platform for tonic and burst firing in nigral DA neurons.

## Results

### Differential contributions of NALCN and TRPC3 channels to tonic and burst firing in nigral DA neurons

We recently reported that NALCN and TRPC3 channels both produce a small leak-like Na^+^ current that induces slow subthreshold depolarization in nigral DA neurons (Um et al., 2021). However, it is not known how these two different ion channels participate in various types of firing in DA neurons. To answer that question, we used the whole-cell patch-clamp technique and recorded the firing activities of nigral DA neurons in midbrain slices from tyrosine hydroxylase (TH)-GFP transgenic mice that express GFP driven by the TH promoter (Figure 1A). As shown in our previous report (Um et al., 2021), selective pharmacological inhibition of either NALCN by L703606 (Hahn et al., 2020) or TRPC3 by pyr10 (Kiyonaka et al., 2009; Schleifer et al., 2012) completely stopped spontaneous tonic firing of DA neurons, and that tonic firing could be reinstated by injecting a leak-like current via a patch-pipette (Figure 1B, C, and D). In the presence of L703606 or pyr10, the rate of revived spontaneous firing correlated with the amount of injected current (Figure 1E), indicating the leak-like properties of the inhibited currents. In both cases, the revived firing rate was restored to the control level by injecting an average depolarizing current of 40 pA, suggesting that NALCN and TRPC3 contribute equally to pacemaking in DA neurons (Um et al., 2021).

**Figure 1.**
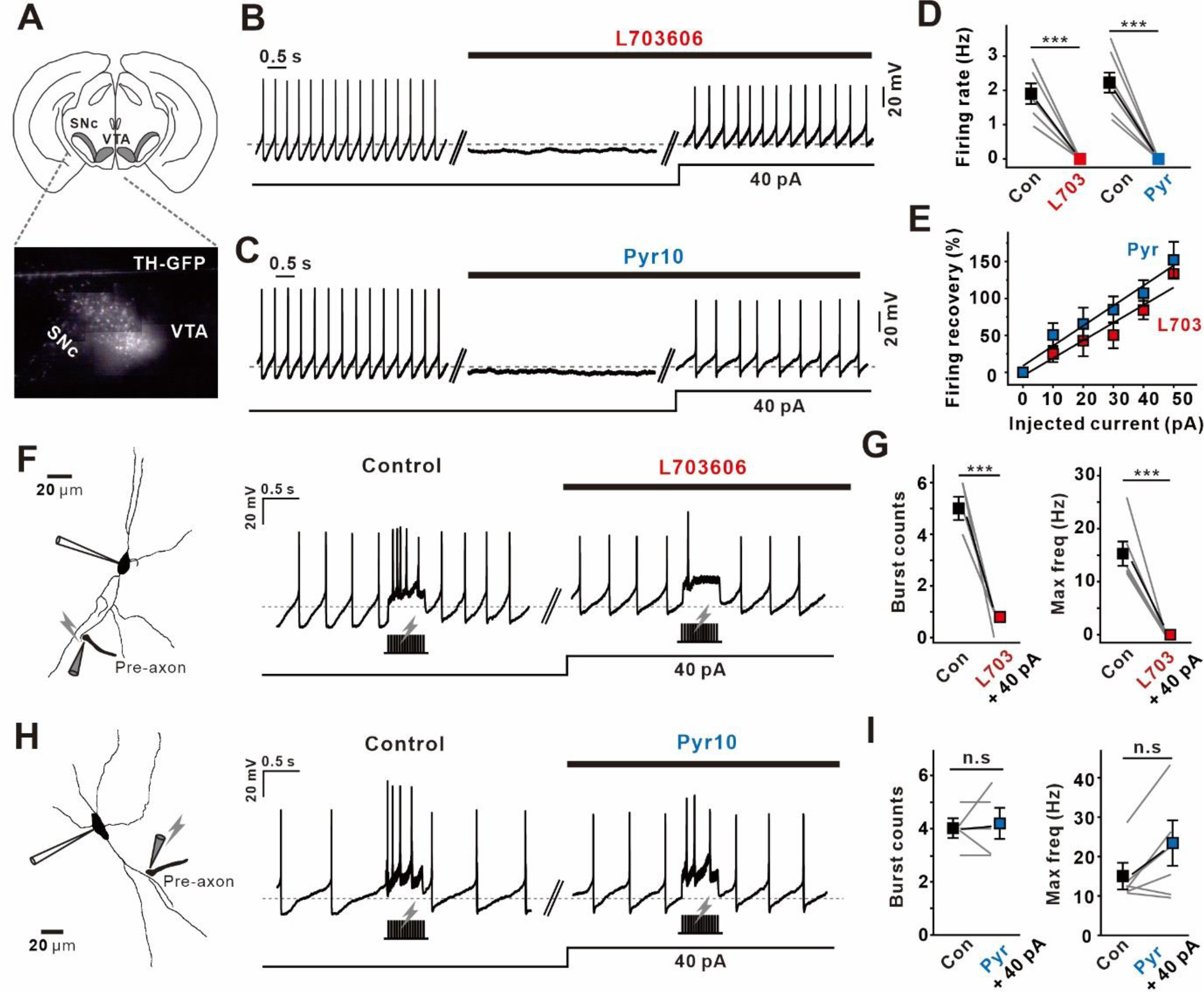
Differential contributions of NALCN and TRPC3 to tonic and burst firing in nigral DA neurons. (A) Schematic diagram of a horizontal section of a midbrain slice with SNc and VTA (upper). Image of SNc and VTA, including GFP-positive TH-DA neurons in a horizontal midbrain slice from TH-GFP transgenic mice (bottom). (B, C) Tonic firing was completely stopped by (B) L703606 and (C) pyr10 application and then rescued by somatic current injections of 40 pA. (D) Summary plots for the effects of L703606 (n=8 from 6 mice) and pyr10 (n=8 from 5 mice) on the tonic firing rate in midbrain slice recordings. (E) Same plots showing the percentages of recovered tonic firing against the injected current sizes in the presence of L703606 (n=22 from 7 mice) and pyr10 (n=25 from 10 mice). (F) 3D reconstruction of a nigral DA neuron from a midbrain slice to visualize the somatic patch clamp pipette and stimulation site (left). The application of L703606 and subsequent somatic current injection recovered tonic firing but did not increase the intensity of burst firing evoked by local electrical stimulation (right). (G) Summary plots of the number of phasic spikes (left) and maximum frequency (right) under the control condition and after treatment with L703606 and subsequent current injection of 40 pA (n=6 from 4 mice). (H) 3D reconstruction of a DA neuron from a SNc slice showing the somatic patch clamp pipette and stimulation site (left). The application of pyr10 and subsequent somatic current injection reinstated both tonic and burst firing to the control level (right). (I) Summary plots of the number of phasic spikes (left) and maximum frequency (right) under the control condition and after treatment with pyr10 and subsequent current injection of 40 pA (n=6 from 5 mice). Closed symbols represent the mean ± SEM. ***p<0.0001, n.s.= not significant (p>0.5). All statistical data were analyzed by the student’s paired t-test.

We next examined whether the NALCN and TRPC3 channels affect high-frequency burst firing in nigral DA neurons. It is well established that local electrical stimulation in midbrain slices evokes burst firing in DA neurons in an NMDAR-dependent manner (Blythe et al., 2007; Hage and Khaliq, 2015). Therefore, we placed a stimulating microelectrode near the dendrites of a patch-clamped DA neuron in midbrain slices (Figure Supplement 1A). Although a weak local electrical stimulus (100 Hz for 0.5 s) slightly affected endogenous tonic firing, the same stimulus given a gradual increase in amplitude eventually elicited phasic firings (Figure Supplement 1A, B, C). Only phasic firings of more than 14 Hz maximum frequency and > 3 spikes were regarded as burst firing in our analyses (Figure Supplement 1B red traces, C right). The recorded burst firing was in accordance with the existing criteria: rapid induction within 100 ms, transient duration (< 200 ms), more than 3 spikes, 14–30 Hz frequency, and interspike intervals (ISIs) between the first and last two spikes of < 80 ms and > 160 ms, respectively (Gonon, 1988; Hyland et al., 2002; Blythe et al., 2007; Blythe et al., 2009). This synaptically evoked burst firing was markedly reduced by the application of 3-(2-carboxypiperazin-4-yl)propyl-1-phosphonic acid (CPP), an NMDAR antagonist (Figure Supplement 1D, E), which is consistent with previous reports (Gonon, 1988; Hyland et al., 2002; Blythe et al., 2007). In our experimental condition, NALCN channel inhibition with L703606 not only eliminated tonic firing but also dramatically attenuated the evoked burst firings (Figure Supplement 2A, C, and D). On the other hand, TRPC3 channel inhibition with pyr10 completely abolished tonic firing, similar to NALCN inhibition, but it did not suppress burst firing (Figure Supplement 2B, C, and D). Because L703606 and pyr10 eradicated tonic firing completely, the disappearance of tonic firing itself might hamper burst generation (Hage and Khaliq, 2015). Therefore, we resuscitated tonic firing by injecting a depolarizing current through a patch-pipette after the application of L703606 or pyr10 (Figure 1F, H). Injecting a 40 pA current into these typical DA neurons restored the tonic firing rate to the control level in both cases. Despite the complete resumption of spontaneous firing, the same electrical stimulation in the presence of L703606 failed to generate burst firing (Figure 1F, G). In contrast, in the presence pyr10, the electrical stimulation evoked burst firing normally (Figure 1H, I). These results suggest that although the NALCN and TRPC3 channels are equally important for tonic firing, only the NALCN channels play a critical role in burst generation in nigral DA neurons.

### Different regional distributions of NALCN and TRPC3 channels in nigral DA neurons

Although NALCN and TRPC3 generate an equal amount of electrically similar Na^+^ leak current in nigral DA neurons (Um et al., 2021), it is unclear how these two ion channels affect burst firing differently. Because the electrical activities of neurons are greatly influenced by the regional distribution of ion channels, we examined the expression patterns of NALCN and TRPC3 in nigral DA neurons using immunofluorescence analyses in the same mice. Because DA neurons in the midbrain extend several dendrites in different directions in a non-linear way, it is very difficult to compare fluorescence intensities across a dendrite in a single plane. Therefore, we used enzymatically isolated single DA neurons with relatively long dendrites attached. Because the dissociated DA neurons attached to cover glass had dendrites spread out in a single plane, we were able to compare the expression patterns of NALCN and TRPC3. Surprisingly, in the DA neurons dissociated from TH-GFP mice, we observed that the intensity of the NALCN immunolabel was highly concentrated in PDCs within 80 μm of the soma (Figure 2A, B, C). Interestingly, the region containing NALCN is identical to that of PDCs < 80 μm from the soma, which we previously reported to be highly excitable and important for generating pacemaker activity in nigral DA neurons (Jang et al., 2014). In contrast, TRPC3 channels were ubiquitously expressed in the whole somatodendritic compartment (Figure 2D, E, F).

**Figure 2.**
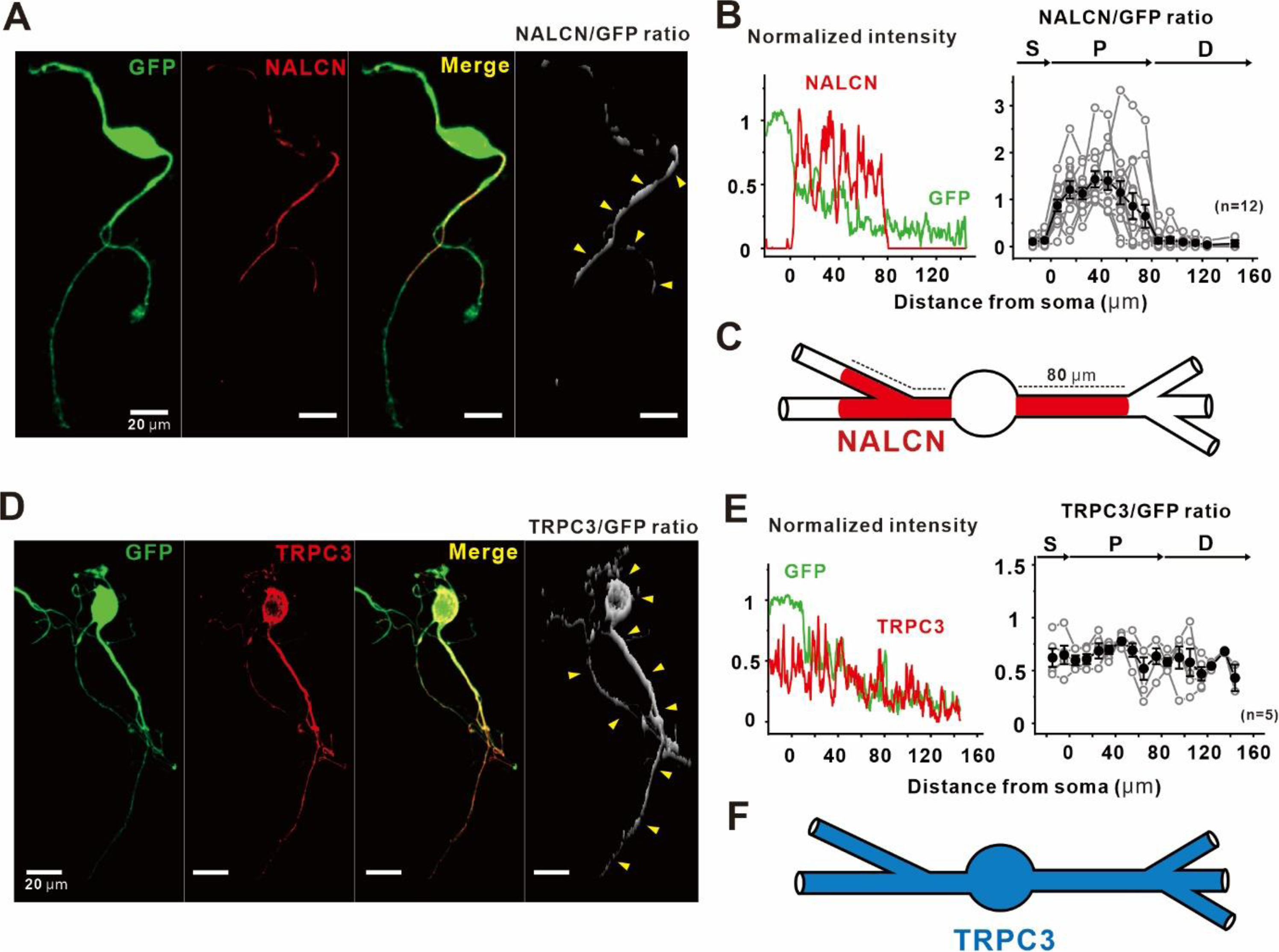
Comparison of the distribution patterns of NALCN and TRPC3 in nigral DA neurons. (A) GFP-positive TH (green, first), NALCN (red, second) labeled by immunofluorescence staining, and merge (third) fluorescence images. The fluorescence intensity ratio of NALCN versus GFP reveals regions of proximal dendrites with concentrated NALCN density (white, fourth). (B) Line profile of the fluorescence intensity of the somatodendritic regions normalized to the fluorescence intensity measured in the soma (left). Plot of the NALCN versus GFP ratio against the averaged binned distance from the soma (right) (n=12 from 6 mice). Peaks of NALCN intensity in the line and fluorescence intensity ratio profiles correspond to the regions of proximal dendrites. (C) Schematic drawing a DA neuron showing the proximal dendritic localization of NALCN, within 80 μm of the soma. (D) GFP-positive TH (green, first), TRPC3 (red, second) labeled by immunofluorescence staining, and merge (third) fluorescence images. The fluorescence intensity ratio of TRPC3 to GFP reveals the ubiquitous expression of TRPC3 in the entire region (white, fourth). (E) Line profile of fluorescence intensity versus distance from the soma, normalized to the fluorescence intensity measured in the soma (left). Plot of the TRPC3 to GFP ratio against the averaged binned distance from the soma (right) (n=5 from 3 mice). No peaks of TRPC3 intensity appear in the line or fluorescence intensity ratio profiles. (F) Schematic drawing of a DA neuron showing the somatodendritic distribution of TRPC3. The yellow arrowheads point to the fluorescence intensity along the somatodendritic regions. Black points represent the average ± SEM. Gray points represent individual data from the averaged binned fluorescence intensity.

### Functional examination of NALCN and TRPC3 channel distributions in nigral DA neurons

To more clearly examine the regional distribution of functional NALCN and TRPC3 channels, we directly recorded NALCN and TRPC3 currents in acutely isolated DA neurons. To activate these ion channels within a localized region of a DA neuron, we used a micro-pressurized puff system (Jang et al., 2014; Hahn et al., 2020). As we previously reported (Hahn et al., 2020), a brief pulse of neurotensin (NT, 10 μM, 1 s puff) induced a substantial amount of NALCN current in dissociated DA neurons, but it was impossible to repetitively activate NALCN currents in a single neuron because of strong desensitization (Hahn et al., 2020). Therefore, we had to activate regional NALCN channels in many different DA neurons and compare the measured NALCN currents (Figure 3A). Consistent with the immunofluorescence staining result, an NT puff on the PDCs evoked a large inward current, but NT stimulation at the soma or distal dendritic compartments did not generate any inward current at all (Figure 3A, B). L703606 completely inhibited the NT-evoked inward currents in the PDCs and also suppressed the basally active NALCN leak currents (Figure 3C, D).

**Figure 3.**
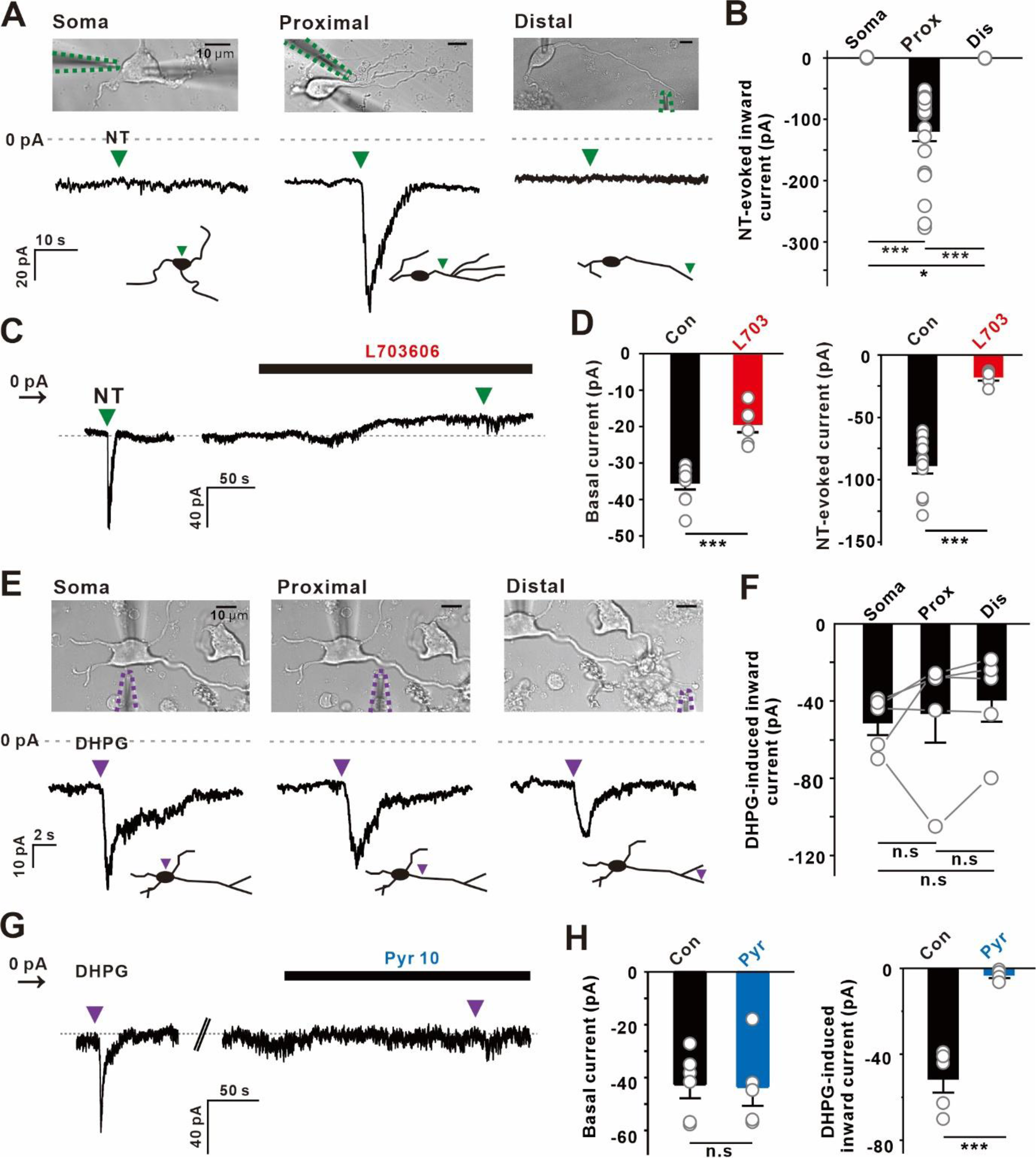
Different generation patterns of NT-activated NALCN currents and DHPG-activated TRPC3 currents. (A) Transmitted images showing the recording pipette and placement of the micro-pressure NT puffing (10 μM, 1 s) pipette (upper, green dotted lines) and the responding current traces with schematic drawings showing the location of each local NT stimulation (bottom). The green inverted triangles indicate NT puffing. Holding potential = −60 mV. (B) Summary of the inward current amplitudes evoked by NT (n=7 from 3 mice for soma, n= 21 from 8 mice for proximal, n=5 from 3 mice for distal). (C) Representative traces of the effect of L703606 (3 μM) on the NT-activated inward current. (D) Summary bar plots of basal currents (left) and NT-evoked inward currents (right) before (n=9 from 4 mice for basal current, n=11 from 5 mice for NT-evoked current) and after the application of L703606 (n=6 from 3 mice for basal current, n=5 from 3 mice for NT-evoked current). Statistical data were analyzed by one-way ANOVA. (E) Transmitted images showing the recording electrode and placement of the DHPG puffing (100 μM, 1 s) pipette (upper) and the responding current traces with schematic drawings showing the location of each local DHPG stimulation (bottom). Example traces were obtained from the same neuron. The purple inverted triangles indicate DHPG puffing. Holding potential = −60 mV. (F) Summary of the inward current amplitudes induced by DHPG (n=5 from 3 mice). (G) Example traces of the effect of pyr10 (3 μM) on the DHPG-activated inward current. (H) Summary bar plots of basal currents (left, n=6 from 4 mice) and DHPG-induced inward currents (right, n=5 from 3 mice) before and after the application of pyr10. Statistical data were analyzed by the student’s paired t-test. Gray dotted lines represent 0 pA. ***p<0.0001, *p<0.05, n.s.= not significant (p>0.5).

However, those results could be attributed to the heterogenous distribution of NT receptors (NTRs), apart from the localization of NALCN channels within PDCs. To exclude that possibility, we performed immunostaining for NT receptors (NTR1). In contrast with NALCN distribution, double immunofluorescence staining showed that TH-positive DA neurons express NTR1 in the entire somatodendritic compartment (Figure Supplement 3A, B). Because NTR1 can also generate IP_3_, which releases Ca^2+^ from the endoplasmic reticulum (ER) Ca^2+^ store (Choi et al., 2003), we checked whether NT induces cytosolic Ca^2+^ increases by using Fluo-4 AM. Local NT puffing to the soma, proximal, and distal dendritic compartments increased [Ca^2+^]c in all cases (Figure Supplement 3C, D, and E). This Ca^2+^ elevation could originate from two distinct sources: Ca^2+^ influx and Ca^2+^ release from the ER. NT-induced Ca^2+^ release from the ER was observed after the removal of all extracellular Ca^2+^ (Figure Supplement 3F). In addition, after emptying the ER Ca^2+^ store with CPA (a sarcoplasmic/endoplasmic ATPase inhibitor), caffeine, an agonist for ryanodine receptors in the ER (Choi et al., 2003; Kim et al., 2004), failed to raise [Ca^2+^]c in DA neurons. Nevertheless, an NT puff on the PDC raised [Ca^2+^]c via Ca^2+^ influx (Figure Supplement 3G). All these data suggest that, while functioning NTR1 is ubiquitously expressed in the whole compartment of a DA neuron, NALCN channels are localized within the PDCs in nigral DA neurons.

Contrary to NALCN, short pulses of dihydroxyphenylglycine (DHPG, 100 μM) were able to repeatedly activate TRPC3 currents (Hartmann et al., 2011) with similar amplitudes in all regions of DA neurons (Figure 3E, F). There was no desensitization to the DHPG-evoked currents in DA neurons, and pyr10 completely suppressed the DHPG-evoked inward currents (Figure 3G, H). All these results are consistent with our immunofluorescence data showing the proximal dendritic localization of NALCN and ubiquitous expression of TRPC3 in DA neurons.

### Importance of highly excitable PDCs in generating the pacemaker activity of nigral DA neurons

Blocking either NALCN or TRPC3 with L703606 or pyr10, respectively, in acutely dissociated DA neurons, as in the midbrain slices (Um et al., 2021), completely abolished spontaneous tonic firing, and the injection of a leak-like depolarizing current revived tonic firing completely (Figure 4A, B, and C). Interestingly, however, after NALCN channel inhibition in acutely dissociated DA neurons, the tonic firing rate was restored to the control level by injecting a 20 pA current, whereas after TRPC3 inhibition, the tonic firing rate was restored to the control level by injecting just a 5 pA current (Figure 4A, B). Thus, under TRPC3 channel inhibition, tonic firing was more easily resuscitated with a smaller current injection than it was after NALCN inhibition (Figure 4F). In the midbrain slices, suppression of tonic firing by TRPC3 or NALCN channel inhibition required a 40 pA depolarizing current to completely recover tonic firing in both cases (Figure 1A–E). Given that the whole-cell NALCN current equals the whole-cell TRPC3 current in intact nigral DA neurons in midbrain slices (Um et al., 2001), the current density of NALCN localized within the PDCs should be higher than that of TRPC3. During the single neuron dissociation procedures, the dissociated DA neurons should lose more distal parts of their dendrites, probably leading to loss of more TRPC3 channels than NALCN channels, which could explain the different results from the current injection experiments in acutely dissociated DA neurons and midbrain slices. Consistent with that interpretation, the input resistance measured in acutely dissociated DA neurons (361.23 ± 12.17, n=7) was about 48% higher than that of DA neurons in midbrain slices (244.44 ± 17.00, n=12). In addition, NALCN channel inhibition with L703,606 increased input resistance significantly more than TRPC3 channel inhibition with pyr10 (Figure 4E). There was no statistical difference between those two groups in the dendritic lengths of the dissociated DA neurons (Figure 4D). All these data suggest that preservation of the highly excitable NALCN-rich PDCs is important for the generation of tonic firing (pacemaker activity) in DA neurons. That interpretation is further supported by the right shift in the firing recovery curve in the current injection experiments after NALCN channel inhibition, compared with TRPC3 inhibition (Figure 4F), unlike the findings in the midbrain slices (Figure 1E).

**Figure 4.**
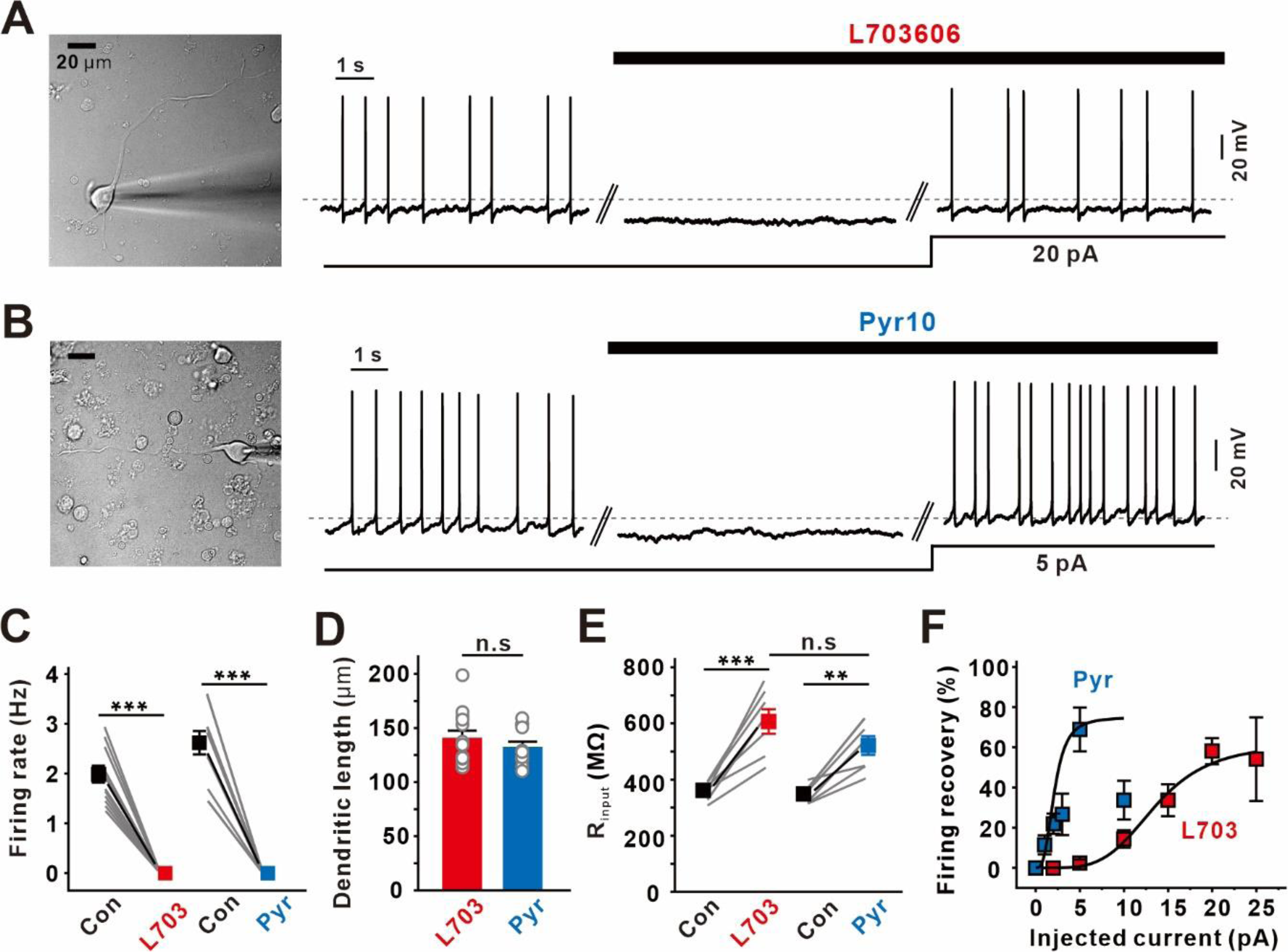
Different depolarizing currents required for the recovery of tonic firing after NALCN and TRPC3 inhibition. (A) Experimental voltage traces of the abolition of tonic firing in the presence of L703606 (3 μM) and the subsequent recovery by current injection in single isolated DA neurons. ***p<0.0001. All statistical data were analyzed by the student’s paired t-test. (B) Voltage traces of the cessation of tonic firing in the presence of pyr10 (3 μM) and the subsequent recovery by current injection. (C) Summary of tonic firing rates before and after treatment with L703606 (n=12 from 4 mice) and pyr10 (n=9 from 4 mice). (D) Comparison graph showing the averaged primary dendritic lengths of DA neurons treated with L703606 (n=13 from 5 mice) and pyr10 (n=12 from 4 mice). Statistical data were analyzed by one-way ANOVA. (E) Summarized graph of the input resistance before and after treatment with L703606 (n=7 from 5 mice) and pyr10 (n=7 from 5 mice). Statistical data comparing the control with the blockers were analyzed by the student’s paired t-test. Statistical data comparing the application of L703606 and pyr10 were analyzed by one-way ANOVA. (F) Different plots showing the percentage of rescued tonic firing against the injected current size in the presence of L703606 (n=34 from 12 mice) and pyr10 (n=54 from 14 mice). Statistical data were fitted by the Boltzmann equation. Gray lines represent individual data. Symbols represent the mean ± SEM. **p<0.01, ***p<0.0001, n.s.= not significant (p>0.5).

These results are in line with our previous report (Jang et al., 2014) that the highly excitable PDCs play a major role in generating the pacemaker activity of nigral DA neurons.

At that time, we demonstrated that a tetrodotoxin (TTX) puff on the PDCs suppressed spontaneous firing more strongly than a puff on the distal dendrites. Taken together, it is likely that the high excitability of the PDCs is due to the localization of NALCN channels.

### Importance of highly excitable PDCs in generating burst firing in nigral DA neurons

Previously, we performed local glutamate uncaging experiments to examine the regional excitabilities of nigral DA neurons (Jang et al., 2014). After silencing DA neurons with a hyperpolarizing current injection, an AP was generated by local glutamate uncaging at a various dendritic sites. As a result, local glutamate uncaging successfully evoked APs only in PDCs < 80 μm from the soma and failed in the distal dendritic compartments, demonstrating that the PDCs had higher excitability than the distal dendritic compartments in DA neurons. Therefore, we examined whether the same highly excitable PDCs also play an important role in burst generation in DA neurons. To generate high-frequency burst firing in dissociated DA neurons, we used the same glutamate uncaging technique as in our previous work. To determine the threshold for generating burst firing in dissociated DA neurons, a single location within the proximal dendritic region was repeatedly stimulated with progressively increasing uncaging areas (Figure Supplement 4A). High-frequency phasic firing with 20–30 μm^2^ uncaging areas corresponds to the definition of burst firing provided earlier (Figure Supplement 4B).

Under those conditions, the same glutamate uncaging area was successively moved along the dendrite of a dissociated DA neuron (Figure 5A). As in the single AP generation experiments (Jang et al., 2014), burst firing was more sensitively generated within the proximal regions (≤ 80 μm from the soma, black circles 1–4) than in the distal parts of the dendrites (black circles 5–6) (Figure 5A). Similar to the previous AP generation experiment with local glutamate uncaging (Jang et al., 2014), burst intensities were induced almost evenly within the proximal dendritic regions, regardless of the uncaging location within the PDCs (Figure 5B). Beyond 80 μm from the soma, the burst generation evoked by local dendritic stimulation disappeared abruptly, as if there were a boundary line. The highly excitable PDCs are also the zone in which NALCN channels are highly expressed (Figure 2C). These data raise the possibility that NALCN localization imparts the high excitability to PDCs that is essential for burst generation (Figure 5C).

**Figure 5.**
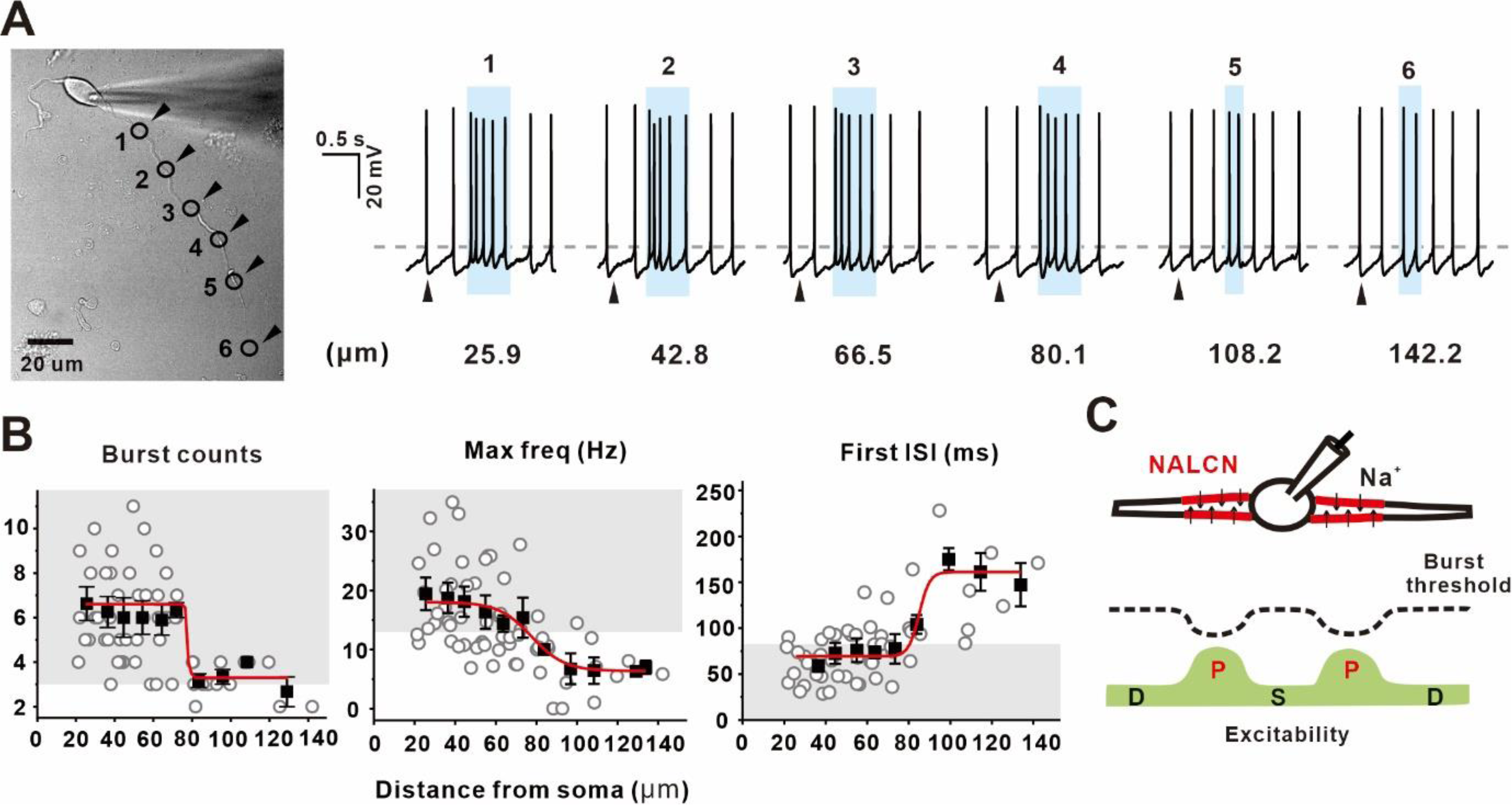
Burst firing is sensitively generated in proximal dendritic compartments of nigral DA neurons. (A) Transmitted image of an isolated DA neuron with the locations of glutamate photolysis (left, black triangles and black circles 1–6) and their corresponding traces of burst firing (right). Burst firing was more sensitively detected with proximal dendritic stimulation than with distal dendritic stimulation. (B) Summarized plots of the number of burst spikes (left, n=60 from 20 mice), maximum frequency (middle, n=61 from 20 mice), and first ISI (right, n=54 from 20 mice) against distance from the soma. All statistical data were fitted by the Boltzmann equation. (C) Schematic drawing of a DA neuron showing high proximal dendritic excitability mediated by the localization of NALCN.

### NALCN localized within the PDCs underlies burst generation in nigral DA neurons

To further examine whether NALCN is directly responsible for burst generation in DA neurons, we evoked burst firing by means of local glutamate uncaging in the proximal dendritic regions of acutely dissociated DA neurons (Figure Supplement 5A–D, control). NALCN channel blockade with L703606 completely suppressed tonic firing. Under that condition, local glutamate uncaging failed to generate burst firing (failure=100%) (Figure Supplement 5A–D, L703606). In contrast, although TRPC3 channel blockade with pyr10 completely stopped tonic firing, local glutamate uncaging produced high-frequency phasic firing in all cases (Figure Supplement 5E–G), and among those cases, more than 60% met the definition of burst firing (Figure Supplement 5F–H).

Therefore, we next investigated whether these different responses between DA neurons silenced by L703606 and pyr10 occurred similarly in normally firing DA neurons. For this experiment, we blocked the NALCN or TRPC3 channels and then restored tonic firing by injecting a leak-like current through a patch pipette in the soma. After that, local glutamate uncaging was performed serially along a dendrite. As shown previously (Figure 5A), local glutamate uncaging evoked burst firing much more effectively in the proximal dendritic regions (< 80 μm from the soma, black circles 1, 2, 3) than in the distal parts of dendrites (> 80 μm from the soma, black circle 4) (Figure 6A, B, control). We then inhibited the NALCN channels with L703606 and restored tonic firing by injecting a leak-like current (average 20 pA). Under that condition, the same glutamate uncaging produced several successive firings, but none of them met the criteria for burst firing (failure=100%) (Figure 6A–C). On the other hand, in the DA neurons whose tonic firing was restored to the control level after TRPC3 channel inhibition, the same glutamate uncaging produced normal burst firing (black circles 1, 2, 3 < 80 μm from the soma, black circle 4 > 80 μm from the soma) (Figure 6E–G). These results are in line with a previous report that tonic firing intensifies burst firing in DA neurons (Hage and Khaliq, 2015). The necessity of resting tonic firing to proper burst firing is further supported by an experiment in which, when DA neurons were progressively hyperpolarized by an increasing current, the same local glutamate uncaging on the PDCs of a nigral DA neuron produced progressively weaker burst firing and finally failed to generate burst firing (Figure Supplement 6). Therefore, although NALCN and TRPC3 generate a similar Na^+^ leak current (Um et al., 2021), it is clear that NALCN channels participate in burst firing differently from TRPC3 channels in DA neurons. The high excitability of PDCs appears to play a critical role in burst generation in DA neurons. The constitutive activities of NALCN channels appear to lower the threshold for burst firing within the PDCs of nigral DA neurons (Figure 6D and 6H).

**Figure 6.**
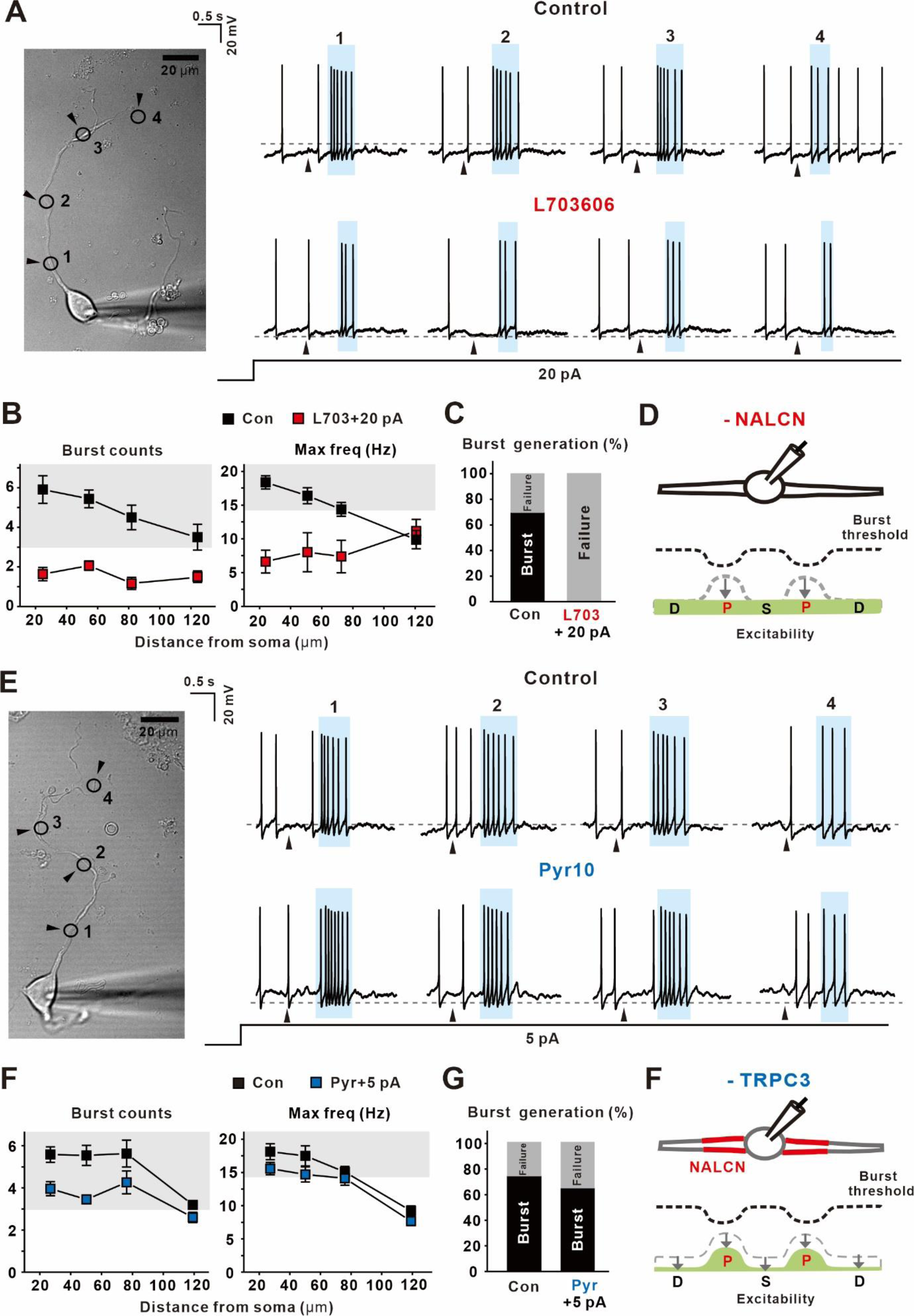
Region-dependent generation of burst firing is sensitive to NALCN blockade but not TRPC3 blockade. (A) Transmitted image of an isolated DA neuron showing the location of each glutamate photolysis (black circles 1–4) (left). Circles 1 (28.31 μm), 2 (54.73 μm), and 3 (75.12 μm) are within the proximal dendritic region, and circle 4 is in the distal region (118.05 μm). Burst intensities were stronger in the proximal dendritic region than in the distal dendritic region under the control condition (right upper). The application of L703606 (3 μM) and an additional current injection of 20 pA reinstated tonic firing but did not increase region-dependent burst intensities at each location of glutamate uncaging (right bottom). (B) Plots of the number of phasic spikes (first, n=53 from 17 mice) and maximum frequency (second, n=49 from 15 mice) against the distance the from soma in the control condition and upon application of L703606 and additional current injections of 20 pA (n=50 from 20 mice). (C) Probability of burst generation before and after the application of L703606 (n=49 from 17 mice). (D) Schematic drawing of a DA neuron with a change in excitability after the application of L703606. (E) Transmitted image of an isolated DA neuron with the location of each glutamate uncaging (black circles 1–4) (left). Circles 1 (18.96 μm), 2 (47.22 μm), and 3 (68.85 μm) are within the proximal dendritic region, and circle 4 (109.32 μm) is in the distal region. Representative traces of the intensities of glutamate-evoked burst firing in the control condition (right upper). Application of pyr10 (3 μM) and a subsequent current injection of 5 pA restored tonic firing and region-dependent burst intensities to near control levels at each location of glutamate uncaging (right bottom). (F) Plots of the number of burst spikes (left, n=64 from 19 mice) and maximum frequency (right, n=55 from 18 mice) against distance from the soma in the control condition and upon application of pyr10 and subsequent current injections of 5 pA (n=55 from 19 mice). (G) Probability of burst generation before and after the application of pyr10 (n=53 from 18 mice). (H) Schematic drawing of a DA neuron showing the change in excitability after the application of pyr10. Black triangles indicate glutamate uncaging stimulation. Light blue boxes represent the duration of burst firing. Gray boxes indicate the ranges corresponding to burst firing. Symbols indicate the mean ± SEM.

## Discussion

In this study, we investigated the cellular mechanisms by which multipolar DA neurons in the midbrain generate tonic and phasic firing. Based on our recent discovery that the TRPC3 and NALCN channels are two major pacemaker channels in midbrain DA neurons (Um et al., 2021), we examined how they participate in tonic and burst firing in nigral DA neurons. We found that, whereas slow tonic firing depends on the basal activity of both the NALCN and TRPC3 channels, burst firing does not require TRPC3 channels but relies only on NALCN channels. In seeking the reason why two functionally similar leak-like channels participate differently in tonic and phasic firing, we found that the TRPC3 and NALCN channels are differently distributed in polarized DA neurons. Whereas TRPC3 is ubiquitously expressed in the entire somatodendritic compartment, NALCN exists only within the PDCs in nigral DA neurons. Given that the whole cell current of NALCN channels is equal to that of TRPC3 (Um et al., 2021), the NALCN channel density localized within the PDCs should be very high, suggesting that NALCN channel localization confers high excitability on PDCs in nigral DA neurons. These results provide strong molecular and cellular evidence for our previous conclusion that PDCs are a highly excitable element that dominantly drives pacemaking in DA neurons (Jang et al., 2014). Therefore, NALCN channel inhibition eliminates the high excitability of PDCs in DA neurons and causes a loss of both tonic and phasic firing. In contrast, TRPC3 inhibition suppresses only tonic firing, but not burst firing. Therefore, it is now clear that the high excitability of multiple PDCs can be ascribed to the basal activities of localized NALCN channels. In addition, we are here the first to demonstrate that multiple PDCs play a key role in not only pacemaking, but also burst generation in nigral DA neurons.

One interesting feature of pacemaking in midbrain DA neurons is strong electrical coupling between the large soma and multiple smaller PDCs (Wilson and Callaway, 2000; Guzman et al., 2009). Therefore, during their pacemaking cycle, DA neurons generate spontaneous APs that are synchronized throughout nearly the entire somatodendritic membrane without significant decrement in their amplitudes (Guzman et al., 2009). In addition, DA neuron pacemaking accompanies Ca^2+^ oscillations, which are also synchronized between proximal and distal dendrites (Guzman et al., 2009). Because DA neurons abundantly express VOCCs (Cardozo and Bean, 1995; Choi et al., 2003) and SK channels (Wolfart et al., 2001; Ji et al., 2009), Ca^2+^ oscillations that occur during the pacemaking cycle cause alternating activation of both VOCCs and SK channels, theoretically affecting membrane potentials. Nevertheless, blockade of those two channels does not seriously jeopardize pacemaking itself (Um et al., 2021; Kim et al., 2007), even though SK channel blockers increase the spontaneous firing rate (Kim et al., 2007). Therefore, it had been suggested that the inward currents produced by VOCC activation might be counterbalanced by an equal amount of outward current produced by SK channels, which are activated by correspondingly increased [Ca^2+^]c (Vrind et al., 2016; Um et al., 2021). Although the tight electrical coupling synchronizes the somatodendritic membrane potential during the slow pacemaking cycle, generating synchronized APs and Ca^2+^ oscillations, [Ca^2+^]c change itself cannot be equal throughout the morphologically heterogenous somatodendritic architecture because the removal time of intracellular Ca^2+^ depends on the surface area/volume ratio of the compartment (Wilson and Callaway, 2000). Ca^2+^ oscillations are greater in the dendrites than in the soma, and they resume more quickly in the dendrites than in the soma (Wilson and Callaway, 2000). Therefore, it has been posited that the rapidly oscillating dendritic compartments are tightly coupled with the slowly oscillating soma, suggesting that dendritic compartments drive pacemaking more than the soma (Wilson and Callaway, 2000). However, the dendritic compartments are not homogeneous in terms of intrinsic electrical properties in most neurons (Magee, 2000). In nigral DA neurons, multiple PDCs appear to be more excitable than the distal dendritic compartments (Jang et al., 2014). Several pieces of evidence indicate that PDCs play a dominant role in pacemaking in nigral DA neurons (Jang et al., 2014). First, our previous morphological and functional analyses showed that the spontaneous firing rate is strongly correlated with the area ratio of the PDCs to the somatic compartment, suggesting that the PDCs are a main player in generating pacemaker activity. Second, inhibiting only the PDCs, but not the distal parts of the dendrites, using a local TTX puff or local Ca^2+^ uncaging slows down the spontaneous firing rates in nigral DA neurons. Third, a computer simulation and local glutamate uncaging experiments reveal that stimulating the PDCs significantly accelerates the pacemaking rate in nigral DA neurons, compared with stimulation of other parts of the neuron. Given that previous evidence of the morphological importance of PDCs for pacemaking in nigral DA neurons, we have collected further evidence about the functional importance of PDCs for pacemaking and burst firing in nigral DA neurons in this study. Most important, we have found that NALCN channels are the molecular reason that PDCs are more excitable than distal dendritic compartments in nigral DA neurons. NALCN channels are spatially localized within multiple PDCs in nigral DA neurons. The PDC regions in which NALCN channels exist (up to 80 μm from the soma) are identical to the more excitable zone for dendrites measured in electrophysiological experiments (Jang et al., 2014). After stopping DA neuron pacemaking by means of TRPC3 or NALCN channel blockade, spontaneous firing recovery experiments with a current injection in acutely dissociated DA neurons revealed the importance of PDCs for both pacemaking and burst firing. Acutely dissociated DA neurons could lose more TRPC3 channels than NALCN channels because they lose more distal dendritic compartments than PDCs during the isolation procedures. Therefore, dissociated DA neurons could have electrical properties that differ from those of intact DA neurons in midbrain slices. Considering the approximately 48% increase in input resistance in our dissociated DA neurons, dissociated DA neurons appear to lose approximately half of their total surface membrane, most likely distal parts of dendrites. Consistent with that measurement, restoring the spontaneous firing rate after TRPC3 or NALCN channel blockade required a smaller current injection in dissociated DA neurons than in intact DA neurons in midbrain slices. Considering the ubiquitous distribution of TRPC3 channels throughout the entire somatodendritic tree and the loss of half the surface membrane, especially distal dendritic compartments, during the dissociation procedure, completely recovering the spontaneous firing rate after TRPC3 channel blockade in acutely dissociated DA neurons required only half the current needed for recovery in intact DA neurons in midbrain slices. Interestingly, the injected current required in the case of a NALCN channel blockade was only a quarter of that required following a TRPC3 channel blockade, indicating that PDCs play a dominant role in pacemaking in nigral DA neurons. The greater increase in input resistance caused by NALCN blockade, compared with TRPC3 blockade, is in line with the results of these firing recovery experiments.

One interesting finding in this study is that PDCs also play a critical role in burst generation in nigral DA neurons. The high excitability of PDCs appears to significantly contribute to the generation of burst firing in nigral DA neurons. Considering the multipolar shape of DA neurons and the many excitatory synaptic inputs throughout the primary dendrites without any fixed spatial input site, the multiple PDCs of slow pacemaker DA neurons would be the best site for integrating various afferent synaptic inputs. For the slow pacemaker activity of DA neurons, multiple highly excitable PDCs and less-excitable soma would be the best choice for both tonic and phasic firing. Therefore, an axon might not arise from the soma because adding a highly excitable AIS to the less excitable soma could cause difficulty in generating high-frequency burst firing. As a result, the proximal dendritic localization of NALCN channels underlies both tonic and burst firings in nigral DA neurons. Therefore, we are now able to propose that multiple PDCs serve as a common platform for tonic and burst firing in nigral DA neurons.

## Materials and methods

### Animals

All experimental procedures were carried out in accordance with the approved animal care guidelines of the Laboratory Animal Research Center in Sungkyunkwan University School of Medicine (Suwon, Korea). STOCK Tg (TH-GFP) DJ76Gsat/Mmnc strain (CrljOri: CD1 background, MMRRC line; RRID: MMRRC_000292-UNC) mice were bred with ICR (Crl: CD1, Orient Bio, Korea, RRID: IMSR_TAC:icr) mice. The mice were housed in sterile cages with no more than five animals per cage and kept in constant conditions of humidity and temperature (21–23℃) and a 12-hour light/dark cycle in a specific pathogen-free vivarium. The mice had access to standard food and water *ad libitum*. Transgenic mice were identified by PCR of genomic DNA obtained from a toe biopsy. The sequences of PCR primers used to amplify the GFP DNA are 5’-CCT ACG GCG TGC AGT GCT TCA GC-3’ (forward) and 5’-CGG CGA GCTGCA CGC TGC GTC CTC-3’ (reverse). We used postnatal day 18–23 mice weighing 9–12 g regardless of sex.

### Slice preparation

Mice were anaesthetized with CO_2_ gas and intracardially perfused with ice-cold high glucose artificial cerebrospinal fluid (aCSF) (in mM: 125 NaCl, 25 NaHCO_3_, 2.5 KCl, 1.25 NaH_2_PO_4_, 0.4 sodium ascorbate, 2 CaCl_2_, 1 MgCl_2_, and 25 glucose, pH 7.3, oxygenated with carbogen gas (95% O_2_/ 5% CO_2_)). After rapid transcardial perfusion, the brains were removed by decapitation and sectioned in 250-μm horizontal slices using a VT-1000s vibratome (Leica, Germany) with high glucose aCSF. The midbrain slices were hemisected and superfused with normal aCSF at 30℃ (mM: 125 NaCl, 25 NaHCO_3_, 2.5 KCl, 1.25 NaH_2_PO_4_, 0.4 sodium ascorbate, 2 CaCl_2_, 1 MgCl_2_, and 10 glucose, pH 7.3, oxygenated with 95% O_2_/5% CO_2_). Slices were then stored in a custom-made maintenance chamber and continuously aerated with carbogen gas until electrophysiological recording.

### Acutely dissociated dopaminergic neuron preparation

The brains were quickly removed in ice-cold, high-glucose HEPES-buffered saline (in mM: 135 NaCl, 5 KCl, 10 HEPES, 1 CaCl_2_, 1 MgCl_2_, and 25 D-glucose, pH adjusted to 7.3 with NaOH (∼310 mOsm/L)) bubbled with 100% O_2_ gas. Coronal midbrain slices were sectioned to 300 μm thicknesses using a TPI vibratome 1000 tissue sectioning system (TPI, USA). The SNc regions were dissected out from each slice and incubated for 20–30 min at 36–37℃ in oxygenated high-glucose HEPES-buffered saline containing 4–8 U/ml papain (Worthington Biochemical Corp., USA, LK0003176). After enzymatic digestion, the SNc pieces were rinsed 3–4 times with papain-free HEPES saline. The tissue pieces were gently triturated using fire-polished pasture pipettes with varying pore sizes to isolate individual cells. The agitated neurons were settled on poly-D-lysine (0.01%)-coated glass coverslips for 30 min at room temperature. All recordings of the single isolated DA neurons were performed within 3 h of dissociation.

### Electrophysiological recording

For brain slice recording, slices were placed into a submersion recording chamber and continuously perfused with fully oxygenated (95% O_2_ /5% CO_2_) aCSF at 33℃. DA neurons were identified and targeted by GFP fluorescence in the SNc regions using an HBO 100 microscope illuminating system (Zeiss, Germany) and a UM-300 CCD camera (UNIQ Vision, USA) on an Axioskop-two microscope equipped with a W plan-apochromat 20× lens (Zeiss). All brain slice recordings were made with an EPC-10 amplifier (HEKA Elektronik, Germany), low-pass filtered at 1–2 kHz, and digitized at a sampling rate of 10–20 kHz with Patch Master software (HEKA Elektronik). To ensure drug delivery to the target neurons, the DA neurons selected for recording were 50– 100 μm beneath the tissue surface (Um et al., 2021).

For whole-cell clamp recordings in the acutely dissociated DA neurons, the cells were perfused with standard external Tyrode’s solution (mM): 140 NaCl, 5 KCl, 10 HEPES, 1 CaCl_2_, 1 MgCl_2_, and 10 D-glucose, pH 7.4 adjusted with NaOH (∼300 mOsm/L) at room temperature. To select GFP-positive DA neurons, the dissociated neurons were exposed to an argon laser (488 nm) and a HeNe laser (633 nm) on a Zeiss 510 confocal microscope. All single isolated DA neuron recordings were performed with an EPC-9 amplifier (HEKA Elektronik, Germany), low-pass filtered at 1 kHz, and digitized at 10 kHz.

The recording pipettes were pulled from filamented borosilicate glass capillaries (World Precision Instruments, USA) on a PC-100 micropipette puller (Narishige, Japan). The patch pipettes had a low-resistance of 4 to 5 MΩ when filled with normal potassium-based internal solutions containing (in mM): 120 K-gluconate, 20 KCl, 10 HEPES, 2 Mg-ATP, 0.3 Na-GTP, 4 Na-ascorbate, 10 Na_2_-phosphocreatine, pH 7.3 adjusted with KOH (280-290 mOsm/L). The liquid junction potentials were −10 mV and corrected for data analysis.

### Local electrical stimulation and imaging

To evoke burst firing, local stimulation of dendritic synapses was performed using unipolar electrodes placed in theta glass pipettes filled with normal aCSF. Electrical stimulation was performed at 100 Hz for 500 ms and placed within 10 μm of a dendrite. Stimulus intensities were set to 10–20 μA. 20 μM SR95531 hydrobromide (Tocris, UK, 1262), 1 μM CGP55845 hydrocholoride (Tocris, UK, 1248), and 1 μM sulpiride (Sigma-Aldrich, USA, S7771) were added to the external solutions to block the GABA_A_ receptor, GABA_B_ receptor, and D_2_ dopamine receptor, respectively.

For fluorescence imaging, DA neurons were loaded with 30 μM Alexa Fluor 594 hydrazide (Thermo Fisher, USA, A10438) via patch pipette and then imaged using a confocal microscope with a 543 nm excitation beam. The laser-scanned images were obtained as Z-sections on an LSM 510 Meta system (Zeiss) and reconstructed to 3D using IMARIS 7.0 (Bitplane, USA).

### Immunocytochemistry

For the immunostaining analyses of NALCN and TRPC3, dissociated DA neurons were fixed with 4% paraformaldehyde (PFA) and 4% sucrose in ice-cold phosphate-buffered saline (PBS, in mM: 137 NaCl, 2.7 KCl, 10 Na_2_HPO_4_, KH_2_PO_4_, pH adjusted to 7.4 with NaOH) for 30 min and then washed with PBS. To stain only TRPC3, fixed neurons were treated with permeabilization buffer (PBS containing 2% normal goat serum (NGS) and 0.1 % Triton X-100) and then rinsed three times with PBS. For NALCN staining, the permeabilization step was omitted. After blocking with blocking buffer (PBS containing 5% NGS) for 45 min, the neurons were incubated in PBS containing rabbit-anti-NALCN (1:50, Alomone Labs, Israel, ASC-022; RRID: AB_11120881) with 2.5% NGS or rabbit-anti-TRPC3 (1:100, Alomone Labs, Israel, ACC-016; RRID: AB_2040236) with 2% NGS overnight at 4 ℃. After the primary antibody reaction, the neurons were rinsed three times with PBS and then incubated with the secondary antibody in PBS for 1.5 hours at room temperature. The secondary antibody was Alexa 647-conjugated goat anti-rabbit IgG (Thermo Fisher, USA, A32733; RRID: AB_2633282) for primary antibody detection of NALCN (1:50) and TRPC3 (1:100). After the secondary antibody reaction, the coverslips were rinsed three times with PBS and then mounted onto slides using fluorescence mount medium (Agilent, USA). Fluorescence images were acquired as Z-sections at 1–1.5 μm × 10 sections with 488 nm (for TH-GFP) and 633 nm (for NALCN or TRPC3) laser lines on a Zeiss 510 confocal laser scanning microscope.

For immunostaining of NTR1, the isolated DA neurons were fixed with 4% PFA and 4% sucrose in ice-cold PBS for 30 min. The fixed neurons were washed three times with PBS and then incubated in permeabilization buffer (PBS containing 2% NGS and 0.1% Triton X-100) at room temperature. After washing them three times with blocking buffer (PBS containing 2% NGS), we administered mouse-anti-NTR1 (1:100, Santa Cruz, USA, sc376958) and rabbit-anti-TH (1:500, Millipore, USA, AB5986P; RRID: AB_92191) to the neurons in blocking buffer for 2 hours at 4℃. Then, the neurons were washed three times with PBS and incubated with secondary antibodies in blocking buffer for 2 hours at room temperature. The secondary antibodies were anti-Alexa-mouse-488 (1:100, Thermo Fisher, USA, A32723; RRID: AB_2633275) and anti-Alexa-rabbit-647 (1:500, Thermo Fisher, USA, A32733; RRID: AB_2633282) for primary antibody detection. After the secondary antibody reaction, the neurons were rinsed three times with blocking buffer and then mounted onto slides using fluorescence mount medium (Agilent, USA). Fluorescence images were acquired as Z-sections at 1 μm × 10 sections with 488 nm (for NTR1) and 633 nm (TH) laser lines on a Zeiss 510 confocal laser scanning microscope. All images were acquired by averaging four images obtained at a resolution of 512 × 512 pixels.

### Micro-pressure injections

For local application of neurotensin (NT) and dihydroxyphenylglycine (DHPG) to each compartment of DA neurons, a pressure microinjection system (Toohey Company, USA) was used. The injection glass pipettes (Model GD-1, Narishige, USA) with resistance between 10 and 20 MΩ were filled with the same external solution containing 10 μM NT (Tocris, UK, 1909) or 100 μM 3,5-DHPG (Tocris, UK, 0805). To prevent the non-targeted compartments from unintended exposure to the agonists, the position of the injection pipette was set within 5 μm of each target compartment. The single puffing duration was 1 s. The injection pressure was maintained between 275 and 345 kPa (Hahn et al., 2020).

### Glutamate uncaging and Ca^2+^ imaging

For local stimulation of glutamate receptors along a dendrite in the dissociated DA neurons, we used 100 μM 4-methoxy-7-nitroindolinyl-caged-L-glutamte (MNI-glutamate, Tocris, UK, 1490) with 10 μM glycine. Uncaging of the MNI glutamate was carried out using a Zeiss 510 confocal microscope equipped with a UV laser (351 and 364 nm) and a HeNe laser (633 nm). The HeNe laser was used to obtain transmitted images during the uncaging experiments. For rapid, localized stimulation, the uncaging speed and power were set to maximum, and the minimum pixel time was less than 1 ms (Jang et al., 2014).

To detect changes in the total cytosolic Ca^2+^ concentration upon local NT stimulation of each compartment, the dissociated DA neurons were incubated with 3 μM Fluo-4 AM (Thermo Fisher, USA, F14201) for 30 min at room temperature. To exclude the effects of the firing activity itself, 0.5 μM TTX was added to the normal external solution. The external Ca^2+^ removal experiments were conducted with Ca^2+^ free external solution (in mM: 140 NaCl, 5 KCl, 10 HEPES, 2 MgCl_2_, and 10 D-glucose, pH 7.4 adjusted with NaOH) and 0.5 μM TTX. In that condition, the puffing pipettes were filled with 0 Ca-based external solution containing 10 μM NT. For depletion of ER Ca^2+^, 10 μM cyclopiazonic acid (CPA, Sigma-Aldrich, USA, C1530) was added to the normal external solution with 0.5 μM TTX (Glentham, UK, GL8460). After the first puff of NT, the dissociated DA neurons were perfused with external solution containing CPA for more than 8 min and then additionally exposed to 20 mM caffeine (Sigma-Aldrich, USA, C0750) to confirm the complete depletion of ER Ca^2+^. The cytosolic Ca^2+^ concentration was measured in all areas of the DA neurons with the maximum scan speed on a Zeiss 510 confocal microscope equipped with a 488 nm laser line.

### Pharmacological reagents

L703606 oxalate salt hydrate (L703606, Glentham, UK, GL7643), pyr10 (Sigma, USA, 648494), CPP (Tocris, USA, 0247), SR95531, CGP55845, sulpiride, neurotensin, 3,5-DHPG, Alexa-594 hydrazide, Fluo-4 AM, TTX, CPA, caffeine, and MNI-caged-L-glutamate were dissolved in dimethyl sulfoxide (DMSO) or deionized water to make stock solutions and then diluted to the final concentration needed for each external or internal solution. All drugs were sonicated before use.

### Data and statistical analysis

All electrophysiological data were analyzed and fitted using Igor Pro 4.01 (Wavemetrics, USA) and Origin 7.0 (OriginLab Corporation, USA) software. All graphical illustrations were made using CorelDraw 2019 software (Corel Corporation, USA). Data were fitted by non-linear fitting with the Hill slope (Figure 4F), Boltzmann slope (Figure 5B), or exponential decay slope (Figure Supplement 1C middle, Figure Supplement 6B left, middle) or by a linear fitting (Figure 1E, Figure Supplement 1C right, Figure Supplement 6B right). Immunofluorescence images were analyzed using Zeiss LSM 510 software. All numeric values and graphical data are presented as the mean ± standard error of the mean (SEM) and number of cells (n). The student’s paired t-test was used to compare two groups, and one-way analysis of variance (ANOVA) was performed to assess the statistical significance of differences among multiple groups. P values of *p<0.05, **p<0.01, and ***p<0.001 were deemed statistically significant.

## Abbreviations

AIS: axon initial segment
AP: action potential
AMPA: α-amino-3-hydroxy-5-methyl-4-isoxazolepropionic acid
CPP: 3-(2-carboxypiperazin-4-yl)propyl-1-phosphonic acid
DA: dopamine
DHPG: dihydroxyphenylglycine
ER: endoplasmic reticulum
GFP: green fluorescence protein
ISI: interspike interval
NALCN: sodium leak channel
NMDA: N-methyl-D-aspartate
NT: neurotensin
PDC: proximal dendritic compartment
SK: channel small-conductance calcium-activated K^+^ channel
SNc: substantia nigra pars compacta
TH: tyrosine hydroxylase
TRPC3: transient receptor potential-canonical 3
VOCCs: voltage-operated calcium channels
VTA: ventral tegmental area

## Acknowledgments

This study was supported by the National Research Foundation of Korea (NRF) grants funded by the Korea government (MSIT) (No. 2017R1A2B3005656 and 2022R1A2C2009159).

## Author Contributions

ConceptuSalization and design, M.K.P., and S.H.; Methodology, analysis and investigation, S.H., and K.U.; Writing – Original draft, S.H.; Writing – Review and editing, S.H., H.J.K. and M.K.P.; Supervision, M.K.P.

## Data availability

All data generated or analyzed during this study are included in the manuscript and have been provided as source data files for all main and supplementary figures.

## Supplementary Figure Legends

**sFigure 1.**
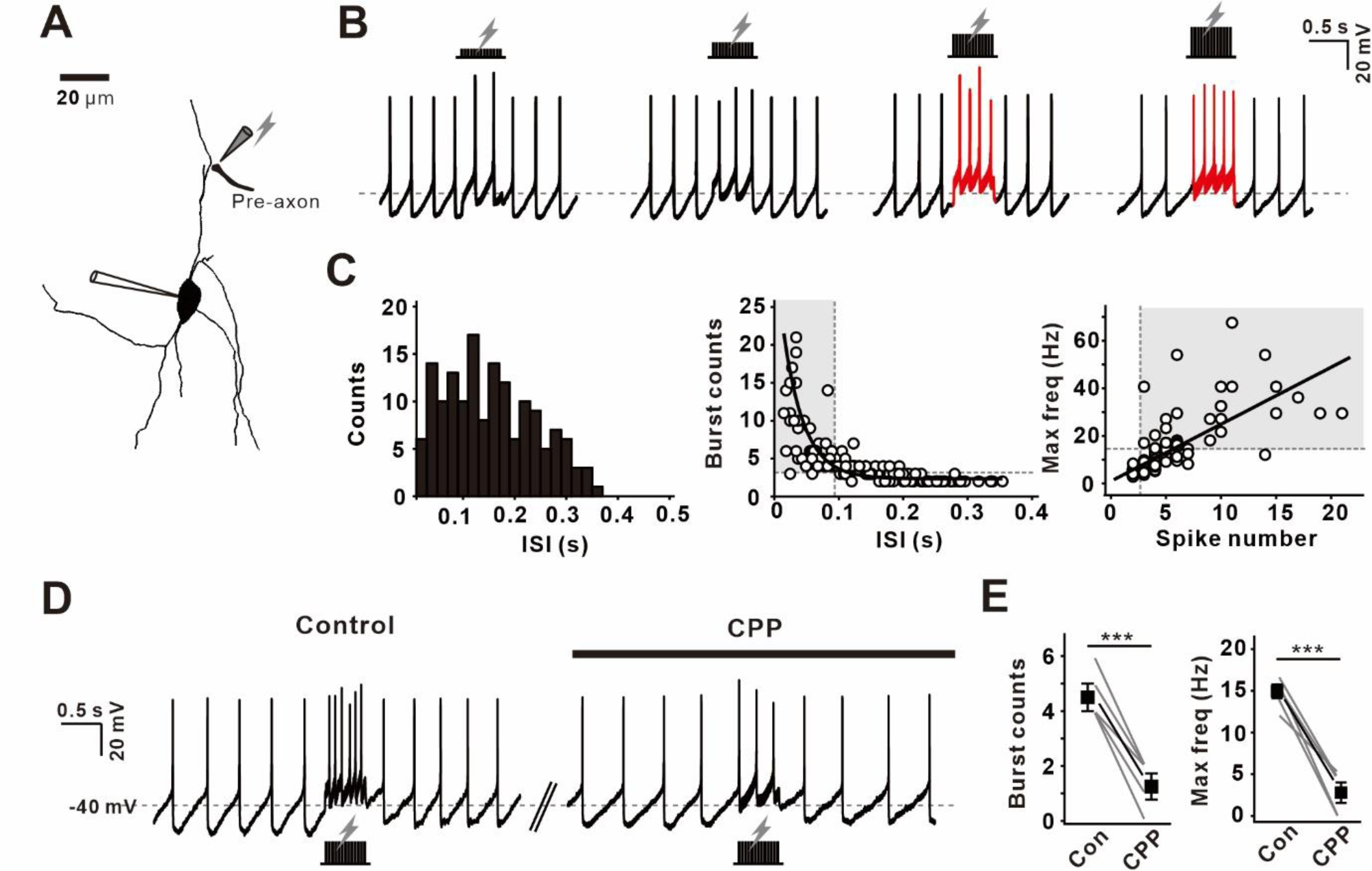
Electrical stimulation evokes NMDAR-dependent burst firing in nigral DA neurons. (A) 3D reconstruction of a TH-GFP DA neuron from a midbrain slice visualized using Alexa-594 through a whole-cell patch pipette. The gray pipette and lightning bolt symbols indicate the stimulation electrode and local stimulation, respectively. (B) Responses of the same neuron (recorded in the whole-cell configuration) to 100 Hz stimulation for 0.5 s with a gradual increase in stimulus intensity. Burst firing is shown in red. (C) All-points histogram of the ISI duration of phasic firing activity evoked by 100 Hz of local stimulation (left), plot of the number of phasic spikes versus ISI duration showing the exponential fit line for all points (middle), and plot of maximum frequency against the number of phasic spikes showing a linear fit for all points (right) (n=150 from 20 mice). The circles are individual points, and the gray dotted lines and boxes represent the bounds and ranges, respectively, corresponding to burst firing. (D) Representative traces in which the intensity of burst firing evoked by 100 Hz electrical stimulation was reduced by CPP. The gray dotted line indicates 40 mV. (E) Summary plots of the number of phasic spikes (left) and the maximum frequency (right) before and after the application of CPP (n=5 from 4 mice). The gray lines represent individual data. Black symbols represent the average ± SEM. ***p<0.0001. All statistical data were analyzed by the student’s paired t-test.

**sFigure 2.**
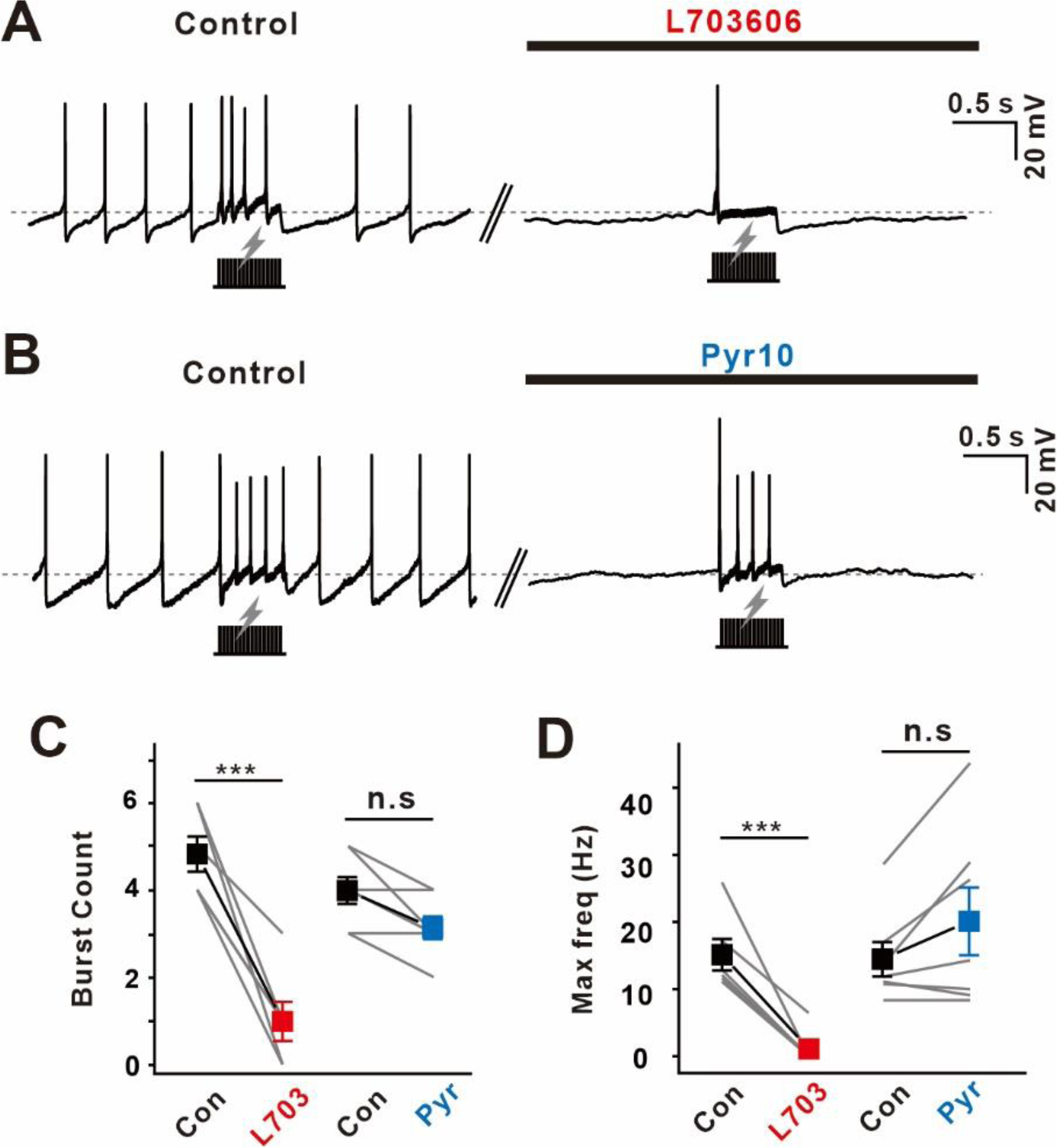
Burst firing is impaired by NALCN blockade but not by TRPC3 blockade after the inhibition of tonic firing. (A) Example traces of burst firing evoked by electrical stimulation before and after the application of L703606 (10 μM). (B) Representative traces of burst firing evoked by electrical stimulation (gray bar) before (control) and after the application of pyr10 (50 μM). (C) Plot of the number of phasic spikes before and after the application of L703606 (n=5 from 4 mice) and pyr10 (n=6 from 4 mice). (D) Plot of the maximum frequency before and after the application of L703606 (n=6 from 4 mice) and pyr10 (n=7 from 4 mice). Black rectangle and gray lightning bolt symbols visualize local electrical stimulation. The closed symbols indicate the average ± SEM. Gray lines indicate individual data. ***p<0.0001, n.s.= not significant (p>0.5). All statistical data were analyzed by the student’s paired t-test.

**sFigure 3.**
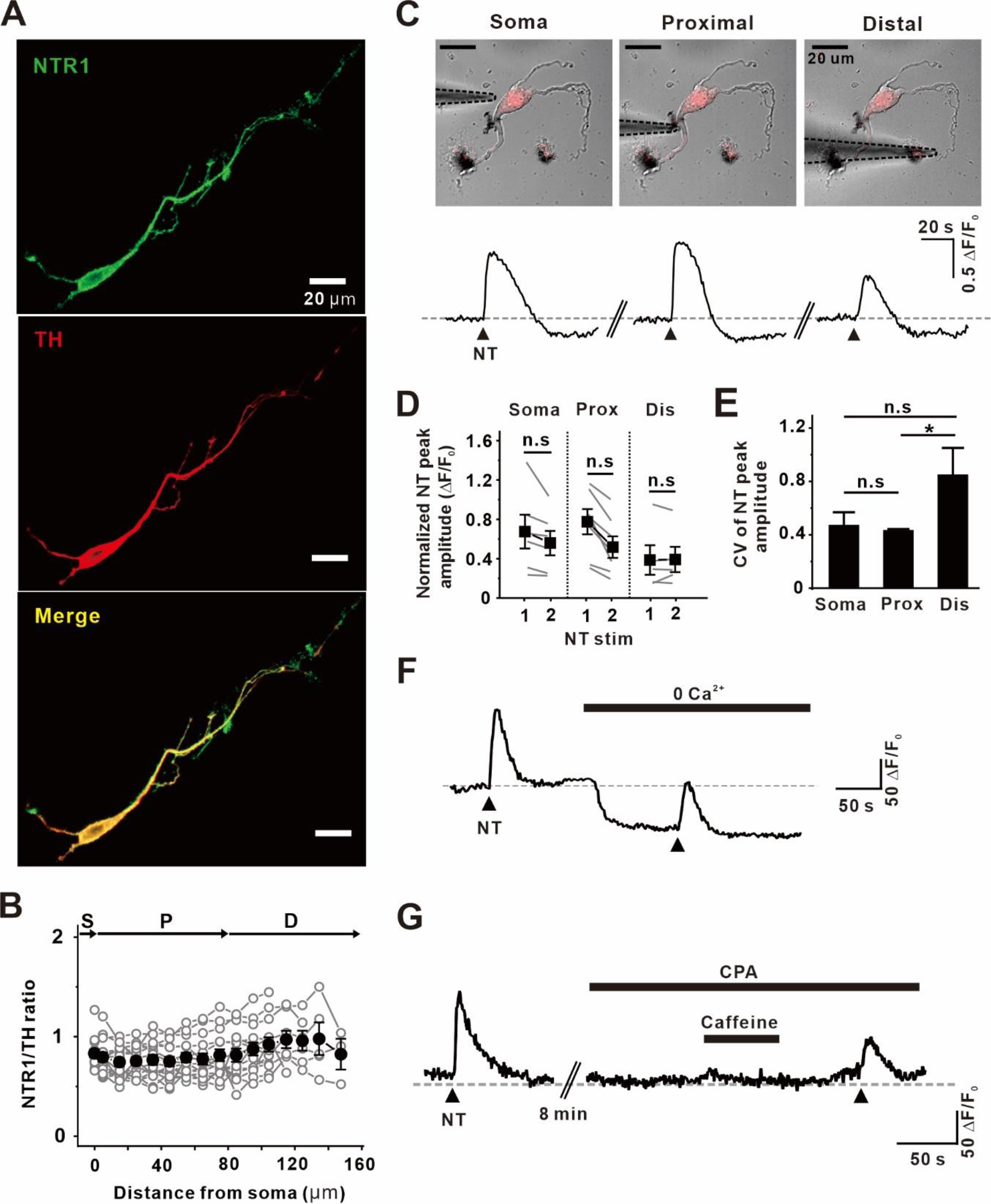
Ubiquitous distribution of NTR1 in nigral DA neurons. (A) NTR (green, top), TH (red, middle), and merge (bottom) double immunofluorescence staining images in a single dissociated DA neuron. (B) Plot of averaged binned NTR1 to GFP ratio against distance from the soma (n=15 from 6 mice). Black points represent the mean ± SEM. Gray points represent individual neurons with averaged binned fluorescence intensity. (C) Transmitted images showing the placement of the NT puffing pipette along the somatodendritic axis and corresponding traces of cytosolic Ca^2+^ imaging (bottom). The black triangles indicate NT puffing. Example traces were obtained from the same neuron. (D) Summarized histogram of the peak amplitude of Ca^2+^ change caused by repetitive NT stimulation of the same area in each compartment (n=6 from 4 mice for soma, n=7 from 4 mice for proximal, n=5 from 4 mice for distal). Repetitive NT puffing repeatedly increased Ca^2+^ in the entire compartment to a similar extent. Cytosolic Ca^2+^ changes were normalized to background fluorescence. Gray lines indicate individual neurons. Black symbols indicate the mean ± SEM. Statistical data were analyzed by the student’s paired t-test. (E) Bar graph of the coefficient variance (CV) of NT-induced Ca^2+^ peak amplitudes in each compartment (n=12 from 4 mice for soma, n=13 from 4 mice for proximal, n=9 from 4 mice for distal). The soma and proximal compartment show less variability than the distal dendritic compartment. Statistical data were analyzed by one-way ANOVA. (F, G) Representative traces of the contributions of (F) Ca^2+^ influx and (G) ER Ca^2+^ release to NT-evoked Ca^2+^ increase. ***p<0.0001, *p<0.05, n.s.= not significant (p>0.5).

**sFigure 4.**
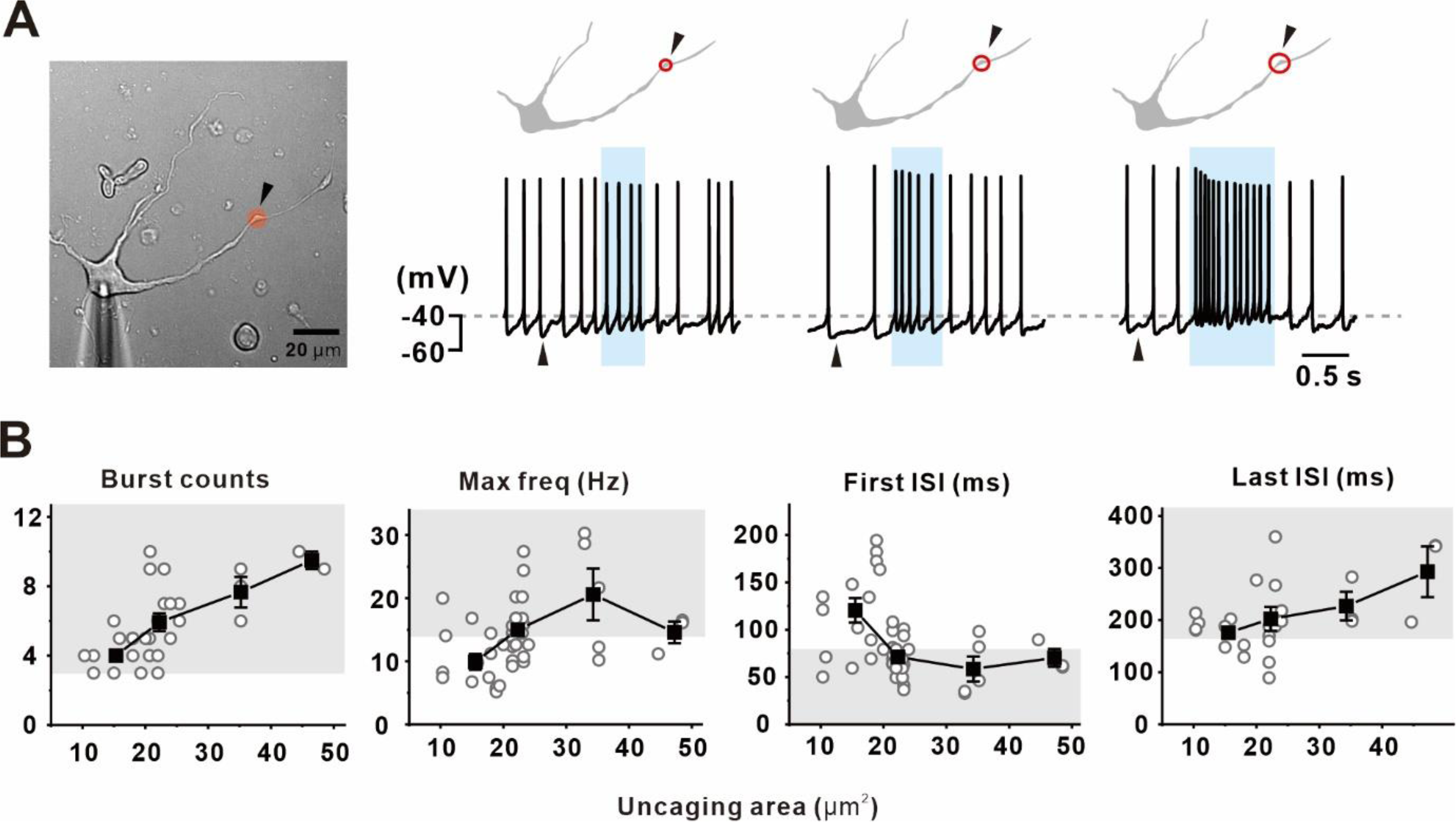
Phasic firing activity correlates with stimulation size of glutamate uncaging. (A) Transmitted image of a single isolated DA neuron with the location of local glutamate photolysis (black triangle and red circle) (left). Representative traces of phasic firing evoked by glutamate uncaging stimulation (right). The intensities of phasic firing gradually increased as the size of the glutamate uncaging area increased. (B) Plots of the number of phasic spikes (first, n=28 from 12 mice), maximum frequency (second, n=46 from 16 mice), first ISI duration (third, n=46 from 16 mice), and last ISI duration (forth, n=26 from 11 mice) against the glutamate uncaging area. Gray points indicate individual data. Black triangles indicate glutamate uncaging stimulations. Black symbols indicate the mean ± SEM. Gray boxes indicate the ranges corresponding to burst firing.

**sFigure 5.**
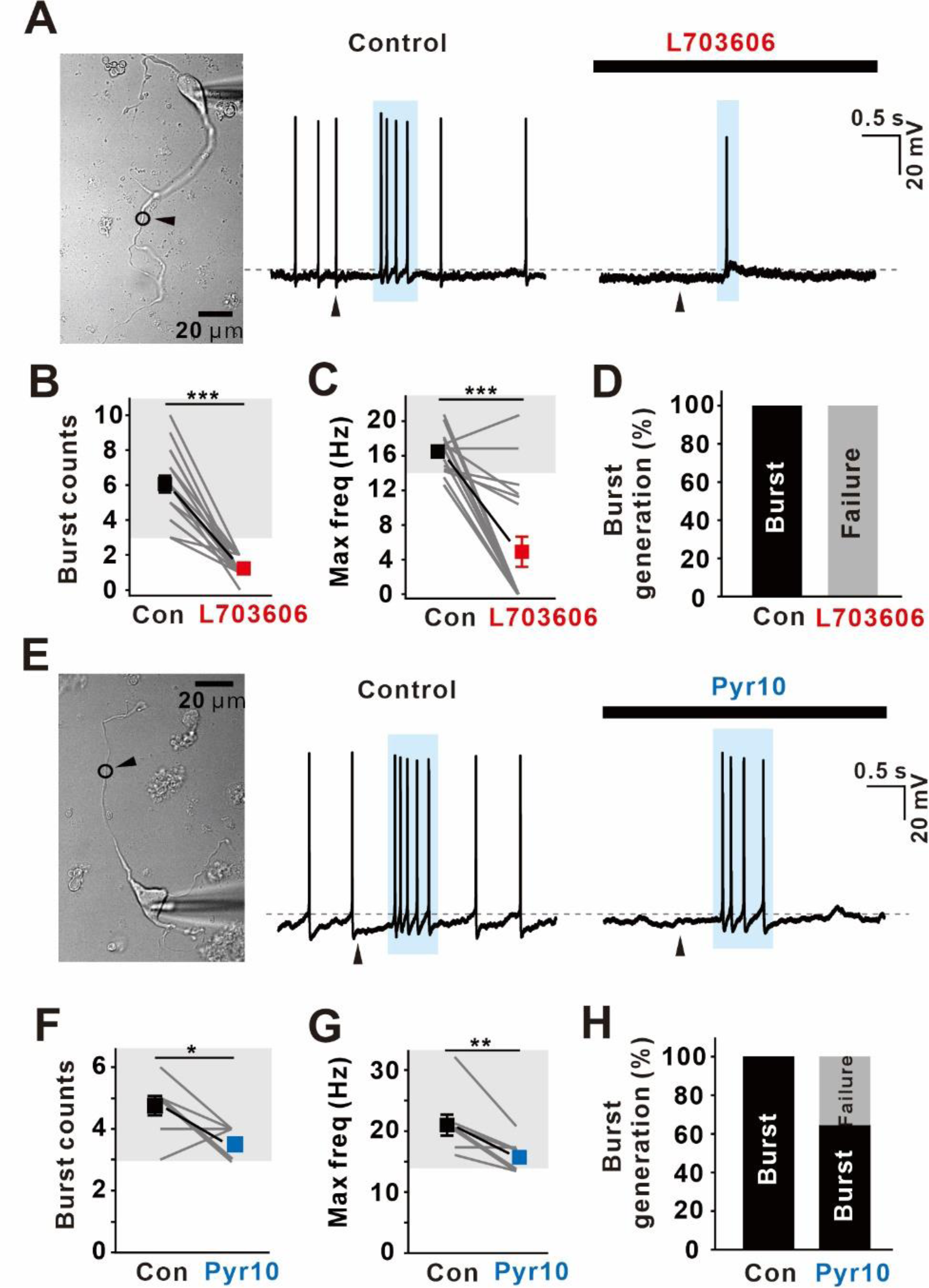
NALCN and TRPC3 blockades differently affect glutamate-evoked burst firing after the inhibition of tonic firing. (A) Transmitted image of a single dissociated DA neuron showing the location of glutamate photolysis (black circle) (left). Glutamate uncaging was stimulated 62.3 μm from the soma. Representative traces of glutamate-evoked burst firing and the effects of L703606 (right). L703606 (3 μM) eliminated both tonic and burst firing. (B) Summary plot of the number of burst spikes before and after the application of L703606 (n=17 from 6 mice). (C) Summary plot of the maximum frequency before and after the application of L703606 (n=17 from 6 mice). (D) The probability of burst generation before and after the application of L703606 (n=25 from 9 mice). (E) Transmitted image of an isolated DA neuron showing the location of glutamate uncaging (black circle) (left). Example traces of glutamate-evoked burst firing and the effects of pyr10 (right). Pyr10 (3 μM) stopped tonic firing but significantly sustained burst intensity. (F) Summary plot of the number of burst spikes before and after the application of pyr10 (n=8 from 4 mice). (G) Summary plot of the maximum frequency before and after the application of pyr10 (n=8 from 4 mice). (H) Probability of burst generation before and after the application of pyr10 (n=17 from 6 mice). Light blue boxes represent the duration of burst firing. Black triangles indicate glutamate uncaging stimulation. Gray boxes indicate the ranges corresponding to burst firing. Symbols indicate the mean ± SEM. *P<0.05, **P<0.01, ***P<0.001. All statistical data were analyzed by the student’s paired t-test.

**sFigure 6.**
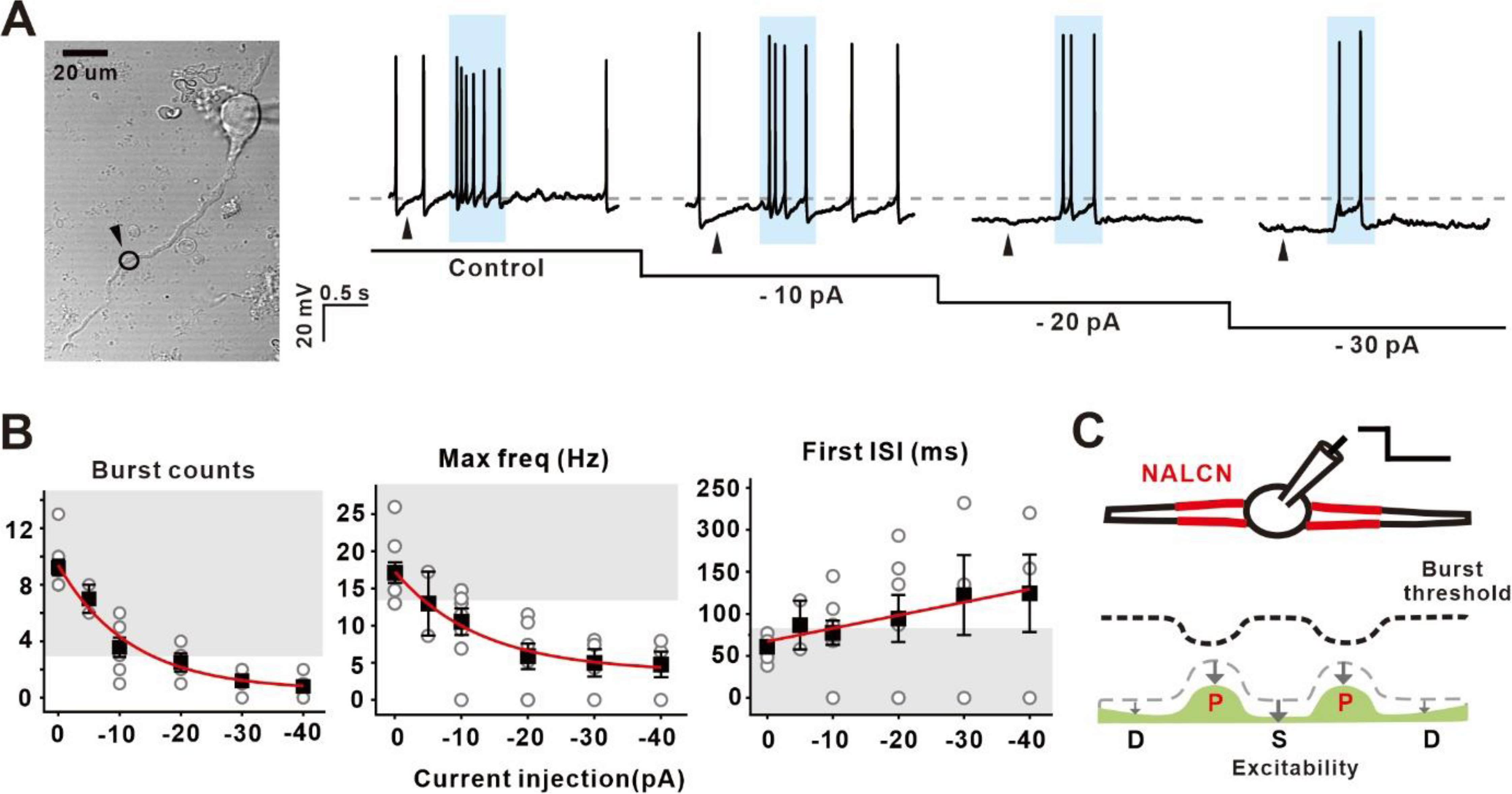
Injection of hyperpolarizing currents attenuates burst firing by decreasing overall excitability. (A) Transmitted image of a single DA neuron with the location of glutamate uncaging in the proximal dendritic region (left, black circle). Example traces of burst firing against hyperpolarizing current injections (right). Burst intensity was attenuated depending on an increase in the injected hyperpolarizing currents. (B) Summary of the number of burst spikes (left), maximum frequency (middle), and first ISI (right) against current injection (n=9 from 4 mice). Statistical data were fitted by an exponential decay curve (burst spikes and maximum frequency) and a linear fit (first ISI). (C) Schematic drawing of a DA neuron whose entire excitability was decreased by an injection of somatic hyperpolarizing current. Black triangles indicate glutamate uncaging stimulation. Gray points indicate individual data. Black symbols indicate the mean ± SEM. Gray boxes indicate the ranges corresponding to burst firing.

